# Harmful algae bloom monitoring via a sustainable, sail-powered mobile platform for in-land and coastal monitoring

**DOI:** 10.1101/473827

**Authors:** Jordon S. Beckler, Ethan Arutunian, Bob Currier, Eric Milbrandt, Scott Duncan

## Abstract

Harmful algae blooms (HAB) in coastal marine environments are increasing in number and duration, pressuring local resource managers to implement mitigation solutions to protect human and ecosystem health. However, insufficient spatial and temporal observations create uninformed management decisions. In order to better detect and map blooms, as well as the environmental conditions responsible for their formation, long-term, unattended observation platforms are desired. In this article, we describe a new cost-efficient, autonomous, mobile platform capable of accepting several sensors that can be used to monitor harmful algae blooms in near real-time. The Navocean autonomous sail-powered surface vehicle is deployable by a single person from shore, capable of waypoint navigation in shallow and deep waters, and powered completely by renewable energy. We present results from three surveys of the Florida Red Tide harmful algae bloom (*Karenia brevis*) of 2017-2018. The vessel made significant progress towards waypoints regardless of wind conditions while underway chl. *a* measurements revealed HAB bloom patches and CDOM and turbidity provided environmental contextual information. While the autonomous sailboat directly adds to our HAB monitoring capabilities, the boat can also help to ground-truth and thus improve satellite monitoring of HABs. Finally, several other pending and future use cases for coastal and inland monitoring are discussed. To our knowledge, this is the first demonstration of a sail-driven vessel used for coastal HAB monitoring.

## 1 Introduction

In the last few decades, harmful algae blooms (HABs) have increased in number, intensity, and duration due to cultural eutrophication, increasing rainfall, and warming temperatures (Brand and Compton, 2007; O’Neil et al., 2012). Through the generation of toxins or by creating locally hypoxic conditions, HAB effects can range from acute sickness and respiratory irritation potentially affecting local economies (Backer et al., 2010; Hoagland et al., 2009; Kirkpatrick et al., 2006), to massive marine fish and mammal mortality events (Gannon et al., 2009; Scholin et al., 2000), or even to chronic human poisoning and death through ingestion of contaminated shellfish or drinking water (Carmichael, 2001; Fleming et al., 2002; Reich et al., 2015). HAB blooms are most frequently observed and anthropogenically detrimental in coastal or in-land marine and freshwater bodies (Anderson et al., 2002), for example in areas with coastal recreation, fishing, mari/aquaculture, and drinking water intake systems. Recent years have experienced superlative HAB events with unparalleled public recognition, for example the summer of 2014 and 2016 Microcystis aeruginosa blue-green cyanoblooms in Lake Erie and the Indian River Lagoon (Florida) (Smith et al., 2015; Stockley et al., 2018) that poisoned drinking water and decreased property values, respectively, the *Pseudo-nitzschia* bloom of 2015 in California waters that led to the closing of the dungeoness crab fishing season (McCabe et al., 2016), and the 2017-2018 *Karenia brevis* bloom in west Florida (ongoing as of the time of writing) that has led to a declaration of a state of emergency. This “Florida Red Tide” bloom is poised to be the worst on record and has brought an unprecedented amount of national attention to this particular HAB (Ducharme, 2018).

To plan for and mitigate the occurrence and effects of HABs, it is ideal to both monitor the algae and/or toxins directly and collect additional ancillary information regarding the chemical and physical ecology of the ecosystems. Traditional routine monitoring is inherently expensive, time consuming, and the spatial and temporal resolution of discrete measurements in many HAB-prone regions is often not sufficient to elucidate bloom causes or properly initiate models. According to a recent HAB scientist community consensus, an observing system consisting of satellite, moored, and mobile data collection platforms will most likely emerge as the most effective holistic approach (Bowers and Smith, 2017). Careful consideration must be given to important tradeoffs existing between sensor specificity targets (e.g. pigments, species, or toxins) and platform compatibility (i.e. fixed location versus mobile), which together determine cost, sampling resolution, and reliability. For example, while satellite-based remote sensing is inexpensive, the technique suffers from insufficient temporal (e.g. daily) and spatial resolution (e.g. ~ 1km), non-species specificity, and interferences from the seafloor, suspended sediment, and clouds. Fixed-location, unattended monitoring devices (i.e. shoreline or moorings) have drastically advanced the temporal resolution of data collection, especially at the species level (Smith et al., 2015; Stockley et al., 2018), but the installation of enough locations to provide sufficient spatial resolution is cost-prohibitive (Shapiro et al., 2015). Given the vertical heterogeneity of HABs, 3-dimensional monitoring platforms, such ocean-going autonomous underwater vehicle buoyancy gliders, are promising and have been successfully deployed in near-shore and open ocean environments (Robbins et al., 2006). However, the submerged nature of these vehicles creates communications, power, and reliability constraints that currently limit sensor options and few species-level options exist. In turn, 2-dimensional Autonomous Surface Vehicles (ASV) such as those powered by sail or waves e.g. “Wave Gliders” or Saildrone (Daniel et al., 2011; Mordy et al., 2017) may alleviate these constraints and are arguably more favorable for more complex instrumentation. However, to our knowledge, all existing long-duration autonomous vehicles are not designed to operate effectively in shallow and/or near-shore waters less than a few meters depth, their size, form or performance prohibit shallow water operation, and their operation is challenging for non-expert resource managers.

For over a decade on the southwest Florida Shelf, fixed location, species-specific optical devices (i.e. Optical Phytoplankton Discriminators; OPD) have been employed as part of a State of Florida and NOAA funded HAB observatory (Sarasota Operations of the Coastal Ocean Observing Lab of Mote Marine Laboratory; SO-COOL). Additionally, AUVs (Slocum gliders) outfitted with either an OPD or a chl. *a* fluorometer are also routinely used to locate and track *K. brevis* HABs (Shapiro et al., 2015). While these efforts have yielded valuable insights into the conditions surrounding HAB bloom formation, these glider operations have presented challenges over the years. Deployments are logistically challenging, requiring an initial transit to deeper waters, and once deployed, a minimum depth limitation of 10 meters (i.e. 20 km from the coast). Finally, cost has prohibited sufficient spatial and temporal coverage, and deployments have been met with unanticipated buoyancy-related operational challenges such as aborts due to nuisance “suckerfish” attaching and sinking gliders (i.e. remora fish).

In 2016, Mote Marine Laboratory began a collaboration with Navocean, Inc., to utilize their autonomous sail-powered surface vehicle for *K. brevis* bloom monitoring. Navocean offers small, 2- m in length vessels that are reliable, and can accept versatile sensors. Navocean boats fill a current niche in both the Autonomous Surface Vehicle (ASV) and the HAB mapping markets, being powered solely from renewable sources, inexpensive, navigable in shallow waters (> 1 m), and deployable from shore by a single person. To demonstrate proof of concept for HAB monitoring, a Navocean *Nav2* boat was outfitted with a 3-channel fluorometer (Turner Designs) configured to measure chl. *a* as a proxy for phytoplankton pigments, as well as CDOM and turbidity to provide ancillary environmental information. The boat was deployed for periods of up to one week in the Winter of 2017, during the start of what has become one of the worst *K. brevis* blooms on record. This work describes the system design, testing, and in situ validation, then discusses other potential applications for HAB monitoring and other environmental applications for this unique vehicle.

## 2 Vessel Design and Operation

The *Nav2* ASV (**Figure 1**) is small, lightweight, easy to launch/land and non-hazardous in the event of collision. The base cost is < $75k and daily operating costs are primarily satellite data fees ($25 to $55 typical). The vessel is 2 m in length, drafts 0.75 m, and weighs between 38 and 45 kg. (depending on battery configuration). The boat has a fiberglass shell with a thick foam core providing reserve buoyancy. The fin keel and rudder are designed to shed seaweed and debris and have proven resistant to tangling in fishing lines and lobster and crab gear in previous missions. The 2 m tall mast has a bright orange sail for high visibility. A “Bermudan” style rig consists of a reinforced carbon mast with high strength Dacron sails (main sail and a small jib) and chafe-resistant lines. The *Nav2* is outfitted with an Airmar 200WX IPX7 marine grade meteorological sensor for wind speed and direction for navigation/scientific purposes, as well as air temperature and barometric pressure for scientific purposes. The *Nav2* is controlled via an iOS application (iPad or iPhone) that is in constant communication to the boat using Wi-Fi, Cellular, or Iridium satellite in either manual mode for line of sight control or autonomous mode for waypoint navigation (**Figure 2**), which includes up-wind tacking in variable wind and sea states. A small electric thruster also provides back-up propulsion for flat calm-wind conditions and for facilitating deployment and recovery, as needed. The standard battery bank consists of up to 5 x LiFePO4 batteries, for a total of 100 A hr and 1200 W hr. Nominal 35 Watt solar panels provide solar recharge of the onboard battery bank for long duration missions (up to several months).

**Figure 1.**
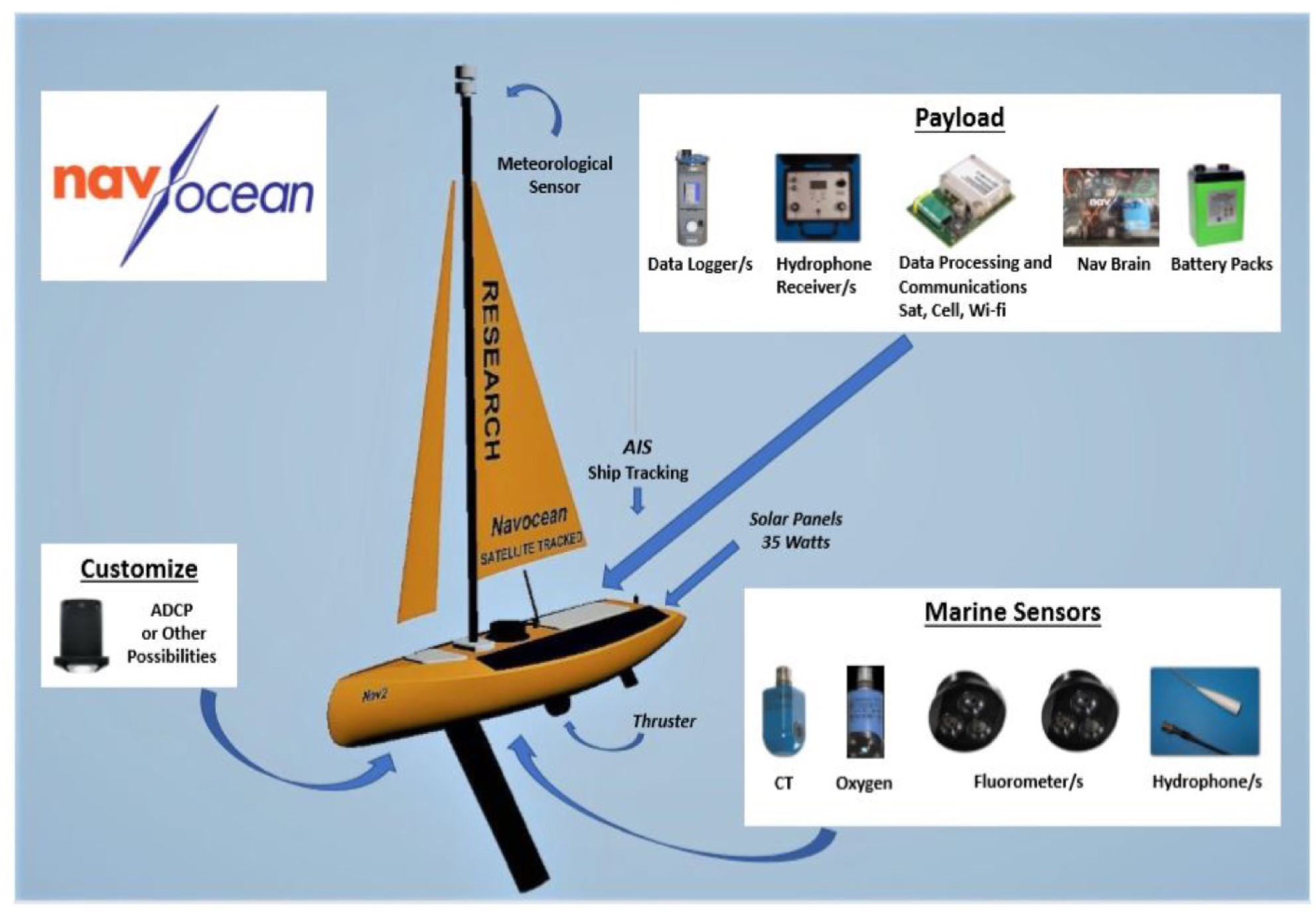
Diagram of the Navocean Nav2 Autononous Sail Vehicle and components.

**Figure 2.**
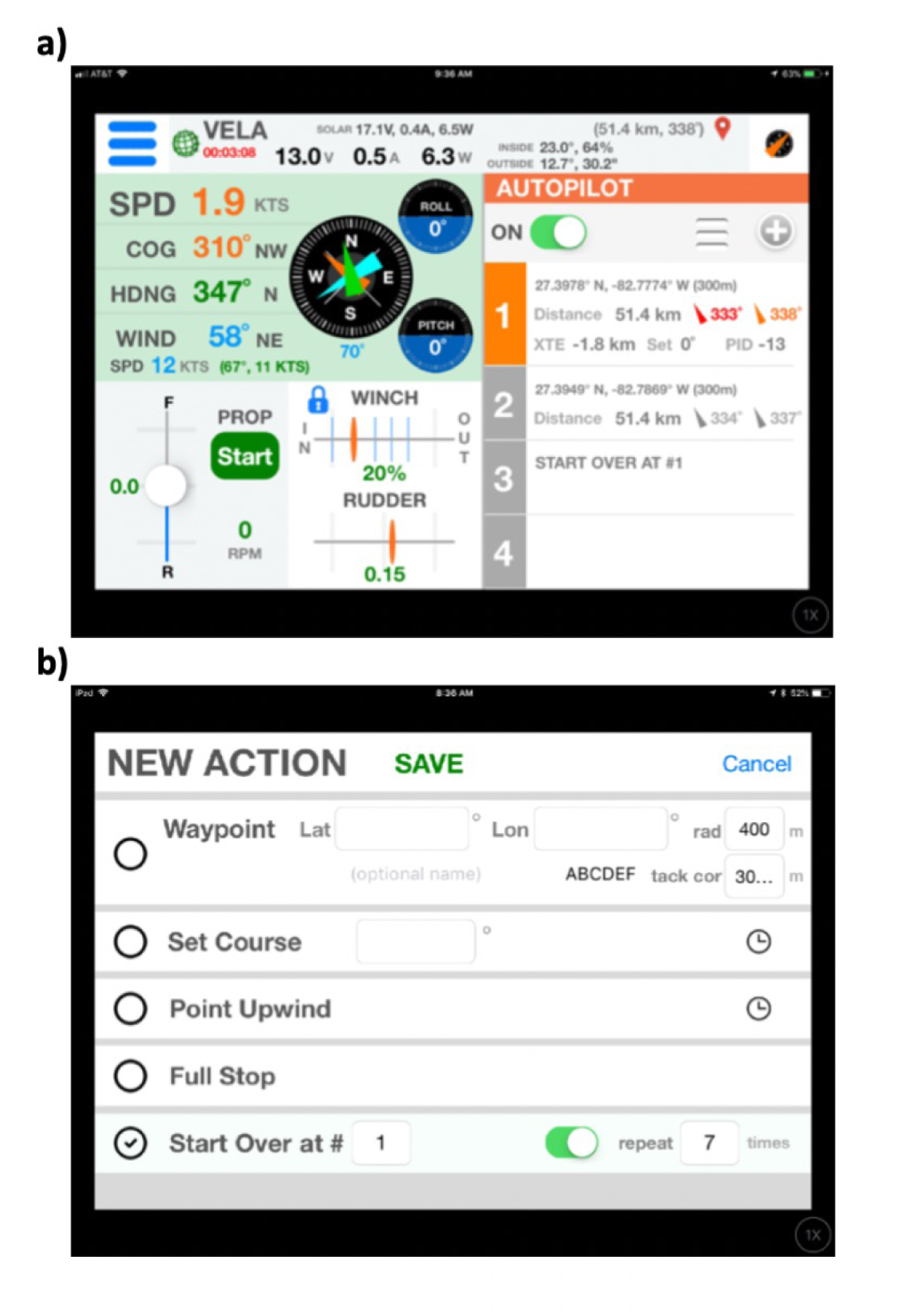
Screenshots of the iOS control software running on an iPad, illustrating operation via Manual Control (**A**) or via Waypoint navigation (**B**).

A 3-channel fluorometer (Turner Designs Cyclops Integrator/C3) configured for measurement of chlorophyll *a*, colored dissolved organic matter (CDOM; measured via fluorescence proxy), and turbidity was installed in the hull, behind the main keel, facing downwards. In all cases, fluorometric measurements are an imperfect measurement and are subject to artifacts. The chl. *a* and turbidity channels underwent single-point cross-calibration using a natural estuarine sample in the laboratory, referenced against a standard benchtop fluorometer that was recently calibrated. The CDOM channel was calibrated instead using the same estuarine water sample, but filtered. The response from the CDOM channel was calibrated using an associated absorption at 440 nm measured in a laboratory spectrophotometer with a 10 cm path length.

**Table 1.**
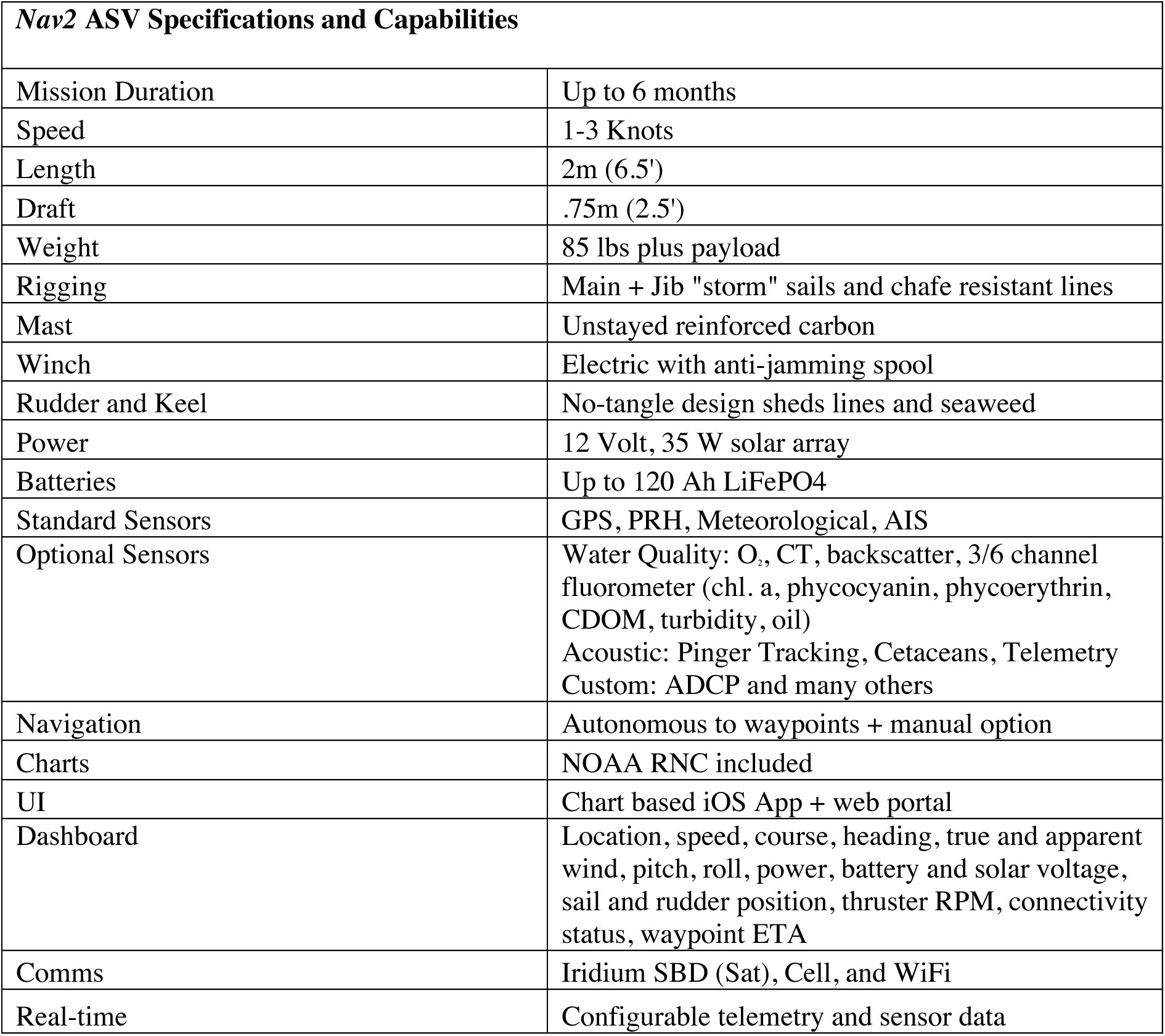
Specifications for the *Nav2* Autonomous Sail Vehicle (Navocean).

| **Nav2 ASV Specifications and Capabilities** |  |
| --- | --- |
| Mission Duration | Up to 6 months |
| Speed | 1-3 Knots |
| Length | 2m (6.5') |
| Draft | .75m (2.5') |
| Weight | 85 lbs plus payload |
| Rigging | Main + Jib "storm" sails and chafe resistant lines |
| Mast | Unstayed reinforced carbon |
| Winch | Electric with anti-jamming spool |
| Rudder and Keel | No-tangle design sheds lines and seaweed |
| Power | 12 Volt, 35 W solar array |
| Batteries | Up to 120 Ah LiFePO4 |
| Standard Sensors | GPS, PRH, Meteorological, AIS |
| Optional Sensors | Water Quality: O<sub>2</sub>, CT, backscatter, 3/6 channel fluorometer (chl. a, phycocyanin, phycoerythrin, CDOM, turbidity, oil)<br>Acoustic: Pinger Tracking, Cetaceans, Telemetry<br>Custom: ADCP and many others |
| Navigation | Autonomous to waypoints + manual option |
| Charts | NOAA RNC included |
| UI | Chart based iOS App + web portal |
| Dashboard | Location, speed, course, heading, true and apparent wind, pitch, roll, power, battery and solar voltage, sail and rudder position, thruster RPM, connectivity status, waypoint ETA |
| Comms | Iridium SBD (Sat), Cell, and WiFi |
| Real-time | Configurable telemetry and sensor data |

## 3 Assessment

### 3.1 Vehicle performance

For the HAB monitoring trials, the *Nav2* vehicle was deployed from the beach three times between Dec. 18, 2017 and Feb. 7, 2018, for deployments of increasing length of one, three, and seven days (**Table 2; Figure 3**), in which case the boat traveled a total of 254 nautical miles (i.e. 470 km; 1 NM = 1.9 km) at an average rate of 1.0 knots (1 knot = 1 NM hr^−1^). Winds were relatively low during this time period, corresponding to an overall average of 6.3 knots as compared to average monthly Dec. and Jan. magnitudes of 10 to 13 knots^1^.

**Table 2.** Summary of the environmental conditions and the *Nav2* ASV performance during harmful algae bloom tracking deployments. The distance covered includes periods of using the thruster at low speeds (~ 1 knot) in calm winds to return the ASV to shore for a convenient pickup time. Alternatively, the thruster can be used temporarily to complete important transects if the wind dies or for entire short missions of ~ 1 day.

| MISSION DATES | NUMBER<br>OF<br>HOURS | WIND<br>SPEED<br>AVERAGE<br>(Apparent) | BOAT<br>SPEED<br>AVERAGE<br>(Knots) | SEA STATE<br>BEAUFORT<br>(Range) | DISTANCE<br>COVERED<br>(NM) | Percent<br>Thruster<br>Use |
| --- | --- | --- | --- | --- | --- | --- |
| Dec 18 to 19, 2017 | 25 | 3.2 | 0.9 | 0-2 | 22.5 | 12% |
| Dec 21 to 24, 2017 | 77 | 3.6 | 0.7 | 0-3 | 53.9 | 10% |
| Jan 31 to Feb 06, 2018 | 148 | 8.2 | 1.2 | 0-5 | 177.6 | 3% |

**Figure 3.**
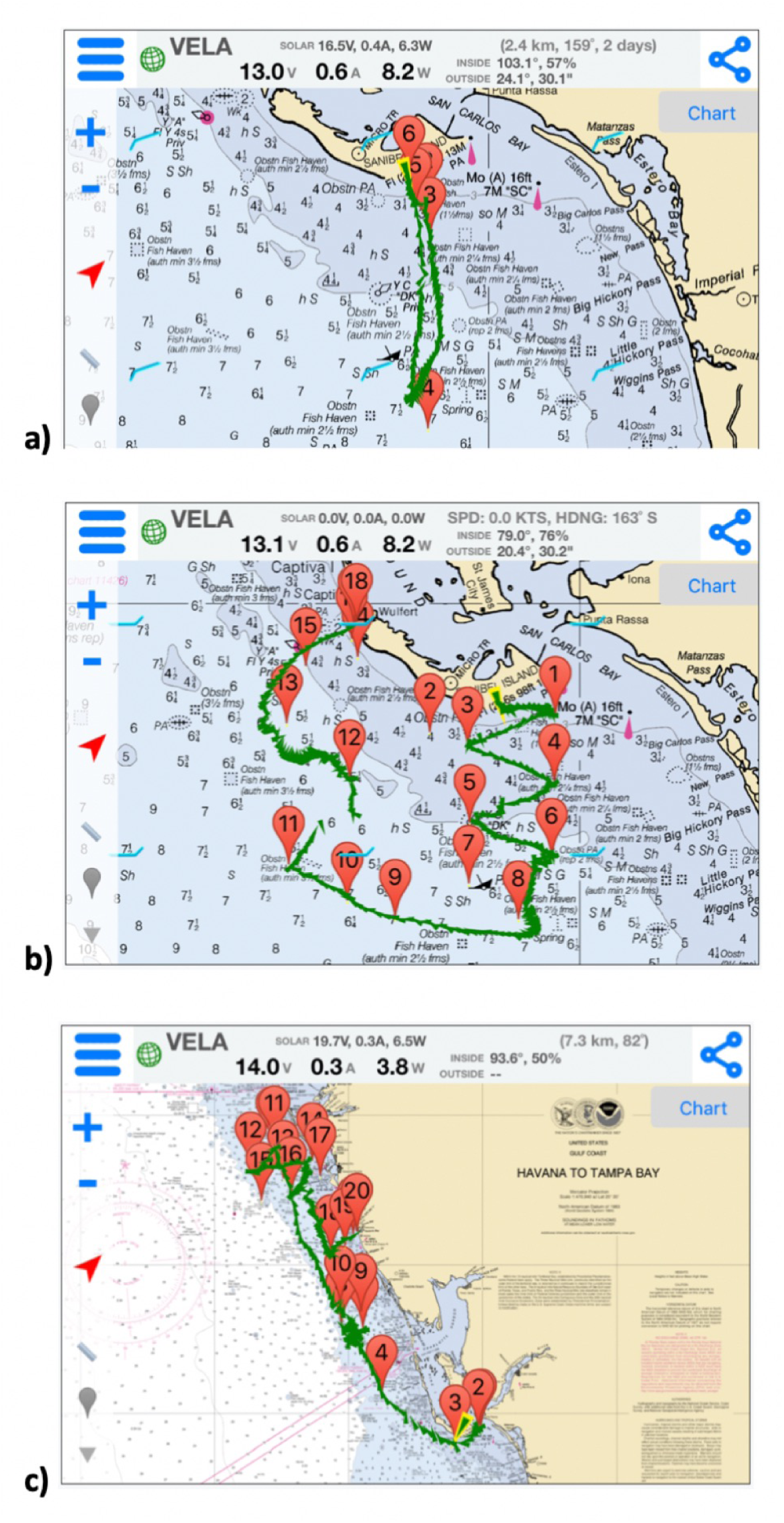
Screenshots of the iOS control software running on an iPad, illustrating the three ASV tracks in southwest Florida for the purposes of harmful algal bloom monitoring from deployments between (**A**) Dec. 18 and 19, 2017, (**B**) Dec. 20 and 23, 2017, and (**C**) Jan. 31 and Feb. 6, 2018. Waypoints

Each mission was operated in a similar manner. The initial waypoints were entered in advance via Wi-Fi using the chart-based app. Iridium satellite communication was used after deployment to monitor the vehicles progress and send updated waypoints as desired, but all navigation was controlled autonomously. To start each mission the *Nav2* was deployed from Sanibel Beach by hand rolling the ASV on its cart out to a depth of > 0.75 m, pointing it offshore, and providing a mild push. At the end of each mission the *Nav2* was directed to sail straight to shore until the keel grounded in shallow water. The *Nav2* was then placed back onto the wheel cart and pulled on-shore. For the 25- hour deployment beginning 2040 UTC Dec. 18, 2017 (**Figure 3a**), the *Nav2* was directed to head straight out and back; sailing first nearly due south to a point 10 NM offshore and then returning north to the beach. On the way back, in response to very calm winds, the thruster was turned on at minimal power to provide a speed of 1 knot which was enough to reach shore at a convenient time for pick up. Some drift was caused by local currents, which presents as a bend in the transect line. Despite the boat experiencing a near full tidal cycle in both the southward and northward direction of travel and experiencing winds between 0 and 3 knots for most of the deployment, the Nav2 steadily progressed. This first short mission served as a data collection test of the fluorometer, which was logging to an SD card on board. For the 77-hour deployment beginning 1422 UTC Dec. 20, 2017 (**Figure 3b**), the *Nav2* was again deployed directly from Sanibel Beach. The intent of this mission was to sail through an area with a known HABs bloom. The Nav2 was directed to first travel south in a zig-zag pattern to cover increased area compared to the first deployment. In response to updated satellite imagery, the *Nav2* was then directed west 15 NM and then north returning to a convenient pick up location at the NE limits of Sanibel Island. The decision was made to persist with sail power for nearly the entire mission to better asses performance in the very calm wind conditions. Depending on solar gain and battery status the thruster can be used for up to 48+ continuous hours to complete straight transects in a timely manner. Tidal current drift effected the precision of transect lines when the wind was < 3 knots. The vessel was removed from the water mid-deployment by a recreational boater, who mistakenly assumed the vessel was lost, and who then traveled with the *Nav2* in a northwest direction for 5 km. The Nav2 was tracked during this time and contact was established with the recreational boaters, who were instructed to place the vessel back into the water. At the end of this mission a more prominent statement was added to the Nav2’s sail indicating boldly its nature as a tracked and monitored research vessel. No such problem has occurred since. Near the end of the mission the winds were calm and the thruster was used at low power to return in a timely manner for pickup at the beach. For the 148-hour deployment beginning1841 UTC Jan. 31, 2018 (**Figure 3c**), the vessel was again deployed from Sanibel Beach with the intent of traversing a significant distance of the West Florida Shelf. The vessel traveled west around Sanibel Island and then proceeded northwards along the coast between 10 and 30 km offshore. After approaching Tampa Bay, the *Nav2* was given waypoints to perform several longitudinal transects, until eventually being directed to the south for retrieval at Venice Beach.

To evaluate the sailing capabilities of the *Nav2* vessel, a polar diagram was constructed (**Figure 4**). The diagram illustrates the obtained vessel speed as a function of realized apparent winds as sensed by the onboard wind sensor (**Figure 1**). For winds from angles directly behind the vessel to as far as 45° into the wind while under waypoint navigation, the vessel autonomously steers directly to the desired destination and the colored lines represent Velocity Made good on Course (VMC). If the Nav2 is traveling towards a desired waypoint that happens to be directly into the wind (with a threshold of 45° port or starboard), the vessel instead autonomously chooses to tack and achieves a net Velocity Made Good (VMG) towards the waypoint. Represented in Figure 4 are therefore two separate calculations; if winds are < 45° off of the bow, the VMG instead represents the apparent velocity with respect to the destination. Increasing apparent wind velocities results in higher *Nav2* velocities for speeds at least as high as 25 knots, under which conditions the vessel is capable of traveling at average speeds >2 knots. The Nav2 is capable of reaching average speeds > 1 knot if winds are at least 5 to 10 knots and greater than 60° away from the wind. Under low wind conditions < 5 knots, the vessel realizes VMC/VMG > 0.5 knots for all apparent wind directions > 30°. Overall, the vessel is capable of realizing significant forward progress, regardless of wind direction, in all but the most unfavorable wind conditions (> 40 km day^−1^).

**Figure 4.**
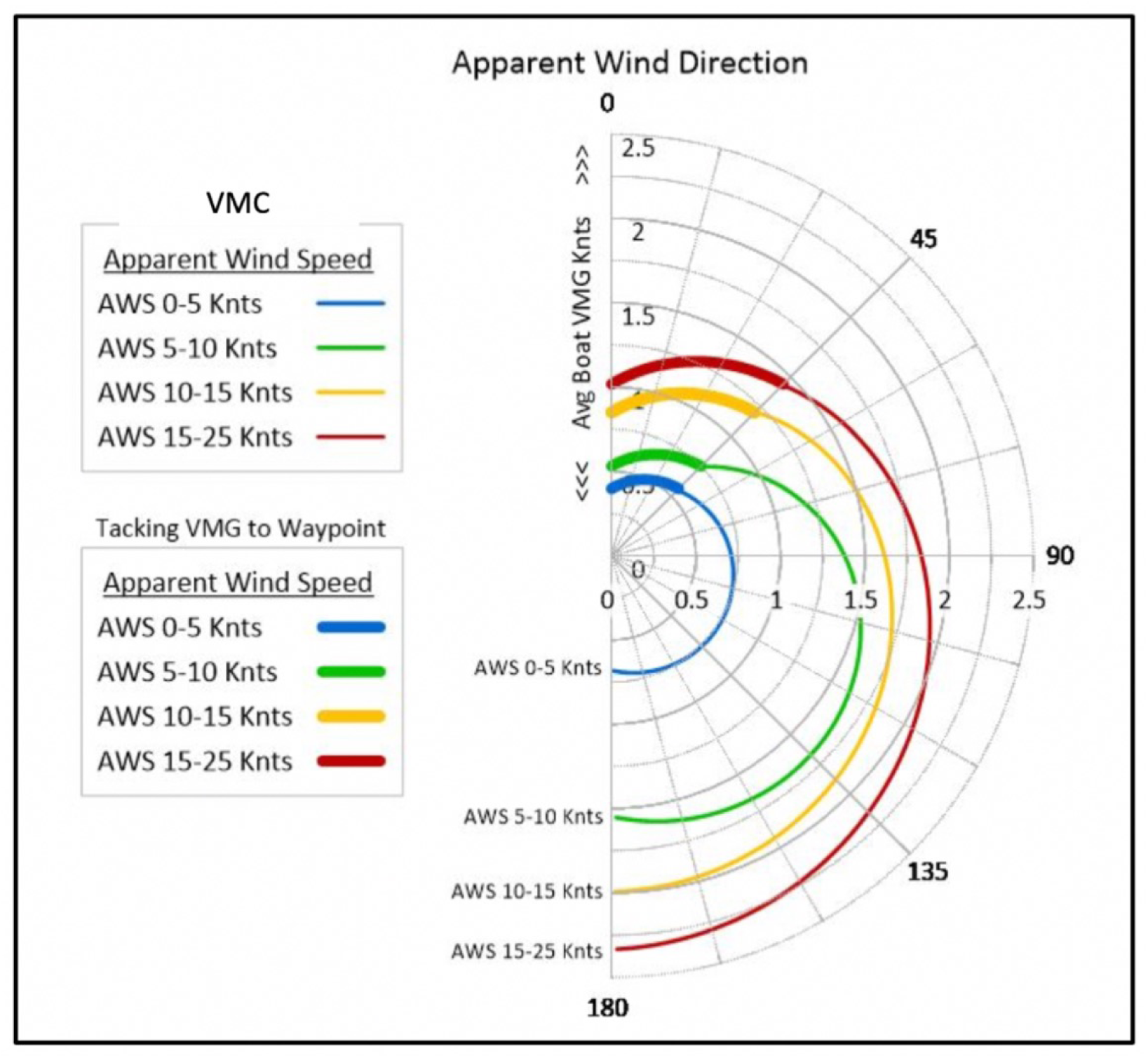
A polar diagram illustrating averaged *Nav2* velocity magnitude while sailing as a function of apparent wind magnitude and direction for all deployments. The four colors represent data for intervals for binned wind speeds. Between angles of 45° and 180°, the magnitude is the actual realized velocity over ground of the vehicle in the intended direction, or the “Velocity Made Good on Course”. The Nav2 tacks as does a traditional sailboat at wind angles < 45°, realizing VMG (“Velocity Made Good”), and the vessel makes significant forward progress even when traveling at very low angles relative to the wind.

To evaluate if there were effects of bubbles on the fluorometric data, the three measured parameters were binned according to the wind speed at the time of data collection (**Figure 5**). We are assuming in this case that higher wind speeds would generate more choppy ocean conditions and thus a larger number of bubbles which may provide measurement artifacts both attenuating and amplifying signals, depending on several factors. However, we observe that the fluorometric data does not appear to depend on wind speed. While there is an increase in chl *a.* values at lower apparent wind speeds, this is likely just coincident with the Nav2 experiencing lower winds closer to shore in the first two deployments, in the presence of the confirmed algae bloom (described in the next section).

**Figure 5.**
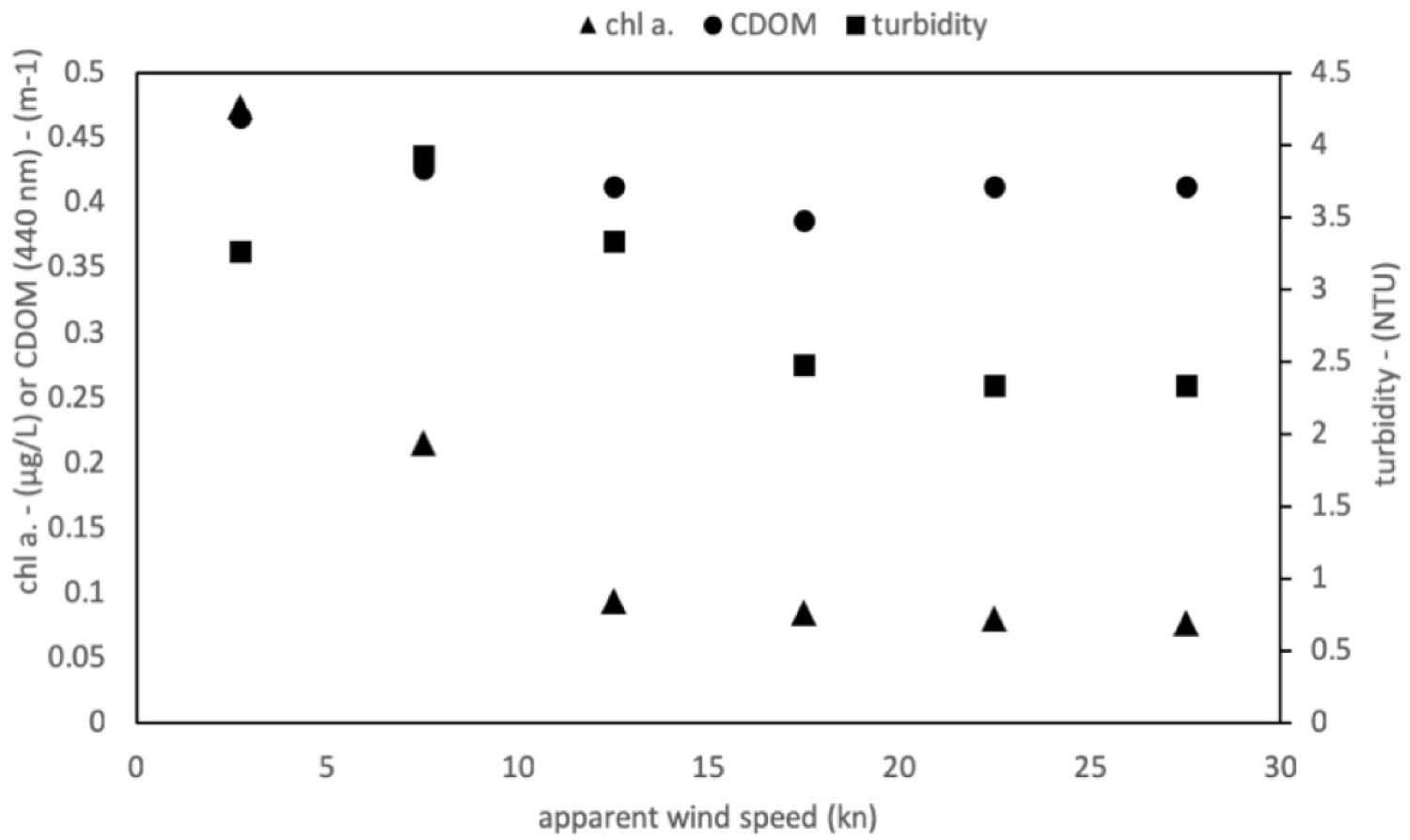
For all deployments, the fluorometer data were binned by their associated 5-knot interval apparent winds speeds to determine if wind and associated bubbles exhibited an artifact.

### 3.2 Harmful algal bloom monitoring

To determine if the Nav2 is a viable platform for HAB detection and mapping, the chl. *a* data was used as a proxy to provide information regarding algal densities. The same 3-channel fluorometer was used as the primary detection means for chl. *a*. For the first two, shorter deployments, a large and intense *K. brevis* (Florida Red Tide) bloom was present near shore (~2 km) towards which the vessel was directed (**Figure 6a&b**). These deployments were intended to demonstrate the potential of the *Nav2* for HAB mapping in localized areas in response to a bloom. The third deployment of 1-week duration on the other hand was intended to demonstrate the potential for the boat to be used for sustained, large-area HAB mapping, even in off-shore environments (**Figure 6c**). Generally, the spatial trends of the in situ fluorometric data agreed well with the results from satellite imagery; however, concentrations derived via remote sensing were significantly elevated compared to the in situ data. For all three deployments, elevated chl. *a* south/southwest of Sanibel Island was probably due primarily to *K. brevis*, given that this species was identified locally at cell counts exceeding 100,000 L^−1^ (discrete samples in **Figure 6a-c**).

**Figure 6.**
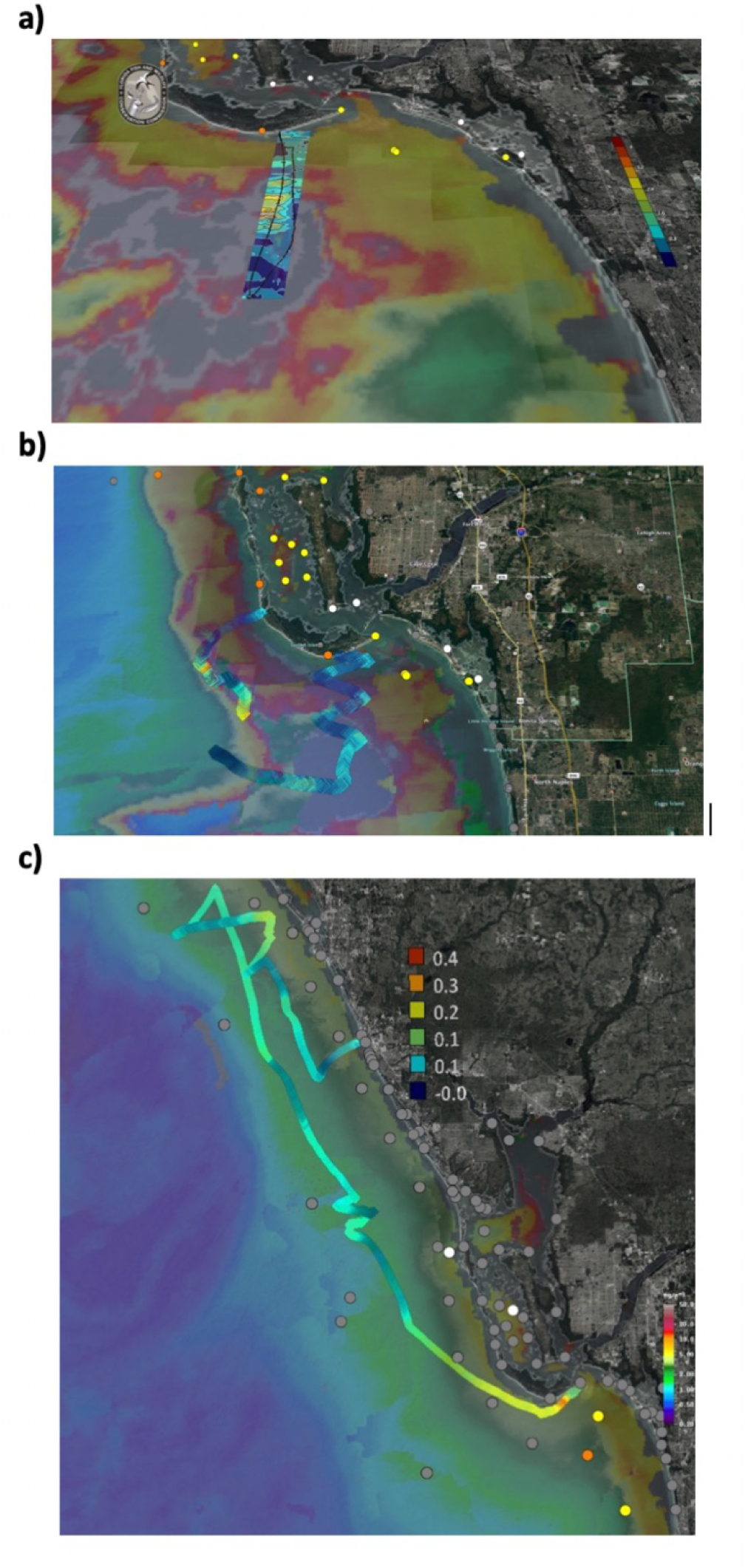
Chl. *a* colormaps or colortracks are presented as demonstrating of HAB mapping capabilities from the three *Nav2* deployments. Background MODIS satellite image is courtesy of Dr. Chuanmin Hu at University of South Florida, and the *K. brevis* cell count data was collected as part of the Florida Fish and Wildlife Commission / Mote Marine Red Tide Monitoring Partnership. The single return raster leg from the deployment during Dec. 18 to 19, 2017 warrants data representation as an interpolated colormap (between 0 and 4 μg L^−1^) with an overlain boat track (black line), (**A**); A colortrack is used to represent (**B**) the Dec. 20-23, 2017 data (between 0 and 4 μg L^−1^), and (**C**) the Jan. 31 – Feb. 6, 2018 data (between 0 and 0.4 μg L^−1^). The background MODIS images are 1-, 3-, and 7-day composites, respectively, and the colors in all represent between 0 and 60 μg L^−1^ remotely-sensed chl. *a* (Note: gray colors indicate areas with cloud cover and no data). Discrete samples collected and enumerated for *K. brevis* cells within 1 week of the deployments are represented by circular icons: grey indicates not present/background levels, white indicates very low densities >1,000-10,000 cells L^−1^, yellow indicates low densities 10,000 – 100,000 cells L^−1^, orange indicates medium densities 100,000 – 1,000,000 cells L^−1^, and red indicates high > 1,000,000 cells L^−1^.

For the first deployment between Dec. 18 and 19, 2017 (**Figure 6a**), the vessel encountered an elevated chl. *a* patch ~2.5 km south of the beach deployment location. Peak in situ concentrations in the patch were ~ 6 μg L^−1^ but were more typically between 1 and 3 μg L^−1^. Interestingly, the initial *Nav2* transect (i.e. southward) only recorded chl. *a* concentrations less than 1.5 μg L^−1^, illustrating the heterogeneity within the patch. In contrast, the remotely sensed background patch was larger and concentrations were higher, between 5 and 50 μg L^−1^. The second deployment between Dec. 20 and 23, 2017 again revealed a high degree of spatial heterogeneity. For the first portion of the deployment chl. *a* concentrations rarely exceeded 2 μg L^−1^. After traveling further west, the vessel soon encountered two chl. *a* patches greater than 5 μg L^−1^. Satellite data did not display a high matchup in this case, as would be expected given that a 3-day composite was used; however, satellite data did reveal that elevated chl. *a* was also observed with a patchy distribution. For the final deployment beginning 5 weeks later between Jan. 31 and Feb. 6, 2018, in situ chl. *a* values were an order of magnitude lower than in Dec. 2017. Just south of Sanibel Island, chl. *a* approached as high as 0.4 μg L^−1^, but then remained less than 0.2 μg L^−1^ for most of the remainder of the deployment. The higher values are consistent with the vessel being closer to shore, but also perhaps with a residual HAB bloom, albeit *K. brevis* cell counts were below detection to the west of the deployment location. While satellite chl. *a* was again much greater than the in situ *Nav2* data, its relative magnitude also decreased by approximately an order of magnitude, with concentrations ~5 μg L^−1^ nearshore and less than 2 μg L^−1^ for the offshore portion of the deployment. Interestingly, several portions of the color track show conspicuously less chl. *a* despite little variations in other parameters, e.g. depth. Upon further investigation, this phenomenon was revealed to be the result of diel variations (**Figure 7c**). Very distinct depressions of the chl. *a* signal were observed between the daylight hours of 1300 and 2300 UTC (8:00 am and 6:00 pm locally). These variations are likely explainable by vertical diel migration (Happey-Wood, 1976) or by variations in pigment expression or measurement artifacts (Babin et al., 1996). These intraday variations were not observed in other deployments where *K. brevis* was likely present (or in the very first day of the 2018 deployment near confirmed *K. brevis*), consistent with the knowledge that this organism does not migrate downwards during the day (Schofield et al., 2006).

**Figure 7.**
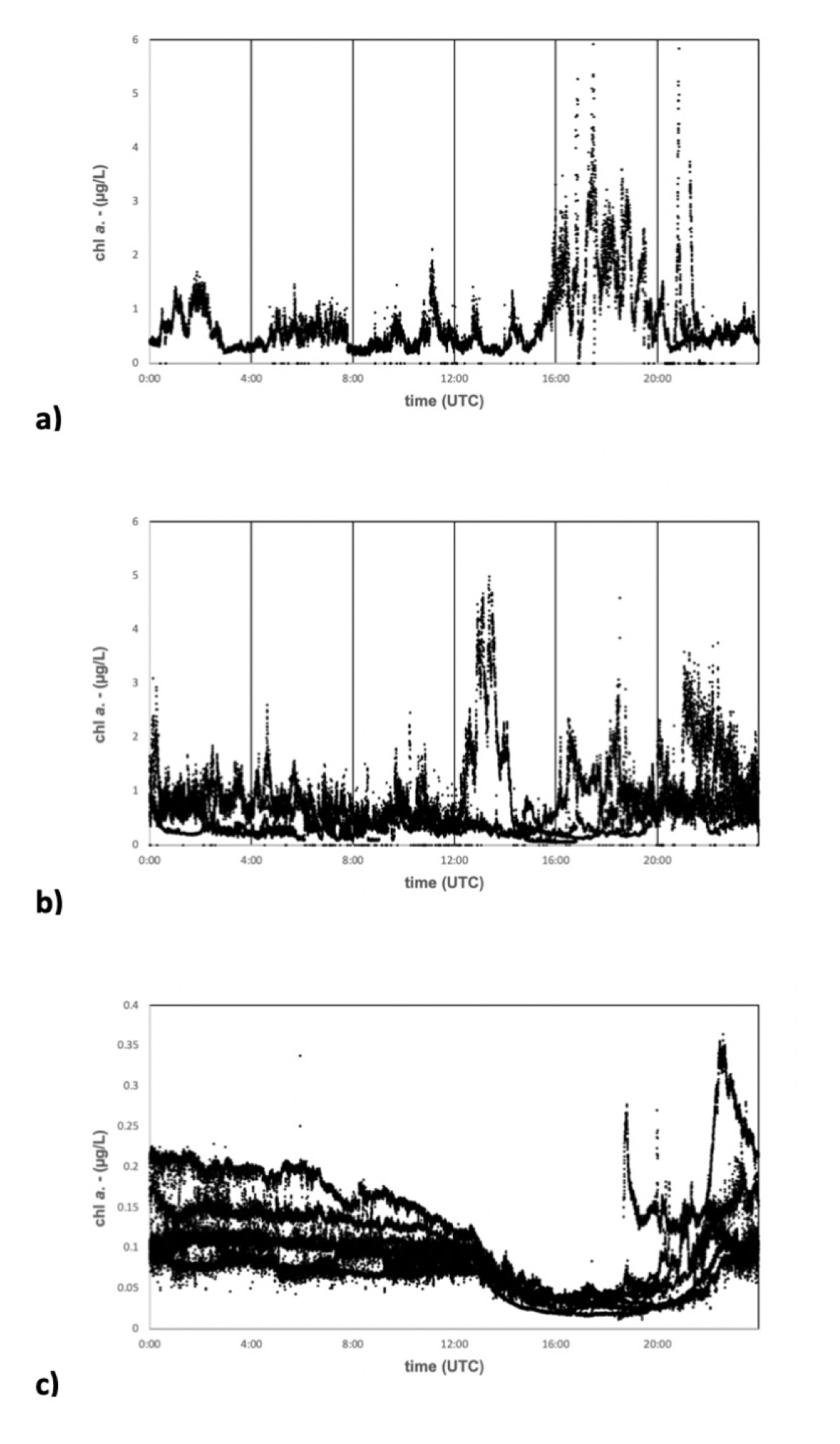
To examine diel trends, daily time series for chl. *a* are presented for the (**A**) Dec. 18-19, 2017, (**B**) Dec. 20-23, 2017, and (**C**) Jan. 31 – Feb. 6, 2018 deployments. Multiple days are depicted on the same plots. Note, the y-axis magnitudes are different for the Jan. 31 to Feb. 6, 2018 deployment.

Turbidity and chl *a*. data provide further information regarding the environmental context of these organisms (**Figure 8d-f**), as well as evidence for the proper functioning of the *Nav2*/fluorometer package, i.e. that the data is consistent with expectations. The CDOM data is represented as absorption at 440 nm despite being fluorometrically obtained. While this is not traditional, we argue that an estimation of CDOM absorption is arguably more useful than representing data in more traditional units (e.g. quinine-sulfate units), and a linear response would be expected either way. Thus, while the CDOM magnitude may not be completely accurate (although values between 0.05 and 0.3 m^−1^ are consistent with CDOM data measured at the Caloosahatchee River outflow)(Del Castillo et al., 2000b)), the spatial variance in the observed CDOM should in fact be accurate. For all three deployments, CDOM increased nearshore consistent with freshwater discharge from inlets, both at deployment and retrieval sites but also during mid-deployment transects (e.g. Feb. 2, 2018; **Figure 8f**). Other increases appear associated with *K. brevis* patches (based on the chl. *a* signature) or river plumes (e.g. Dec. 19, 2017; **Figure 8d**). Along these lines, in the absence of a bloom and in a coastline receiving discharge from a single freshwater source, the CDOM data may serve as a proxy for salinity. Turbidity, being measured as the amount of light scattered at 90° from a source at a single wavelength, appears to more reflect a combination of suspended sediment and phytoplankton cells (**Figure 8g-i**). Turbidity measurements were more transient and less precise at a single location than CDOM (e.g. Feb 3,4, 2018; **Figure 8i**), consistent with transient suspended sediments and a heterogeneous water column. Winds were indeed in the 12 to 25 knot range Feb 3 from around UTC 0600 to 2200, and then perodically elevated on Feb. 4 throughout the day, which could provide an explanation. Turbidity increases were also observed near *K. brevis* bloom patches (evidenced by elevated chl. *a*; **Figure 8g**). It is notable that CDOM measurements have a higher precision than the turbidity measurements (e.g. **Figure 8f&i**). This is expected because the dissolved CDOM will be much more homogenously mixed than will particulates measured via turbidity.

**Figure 8.**
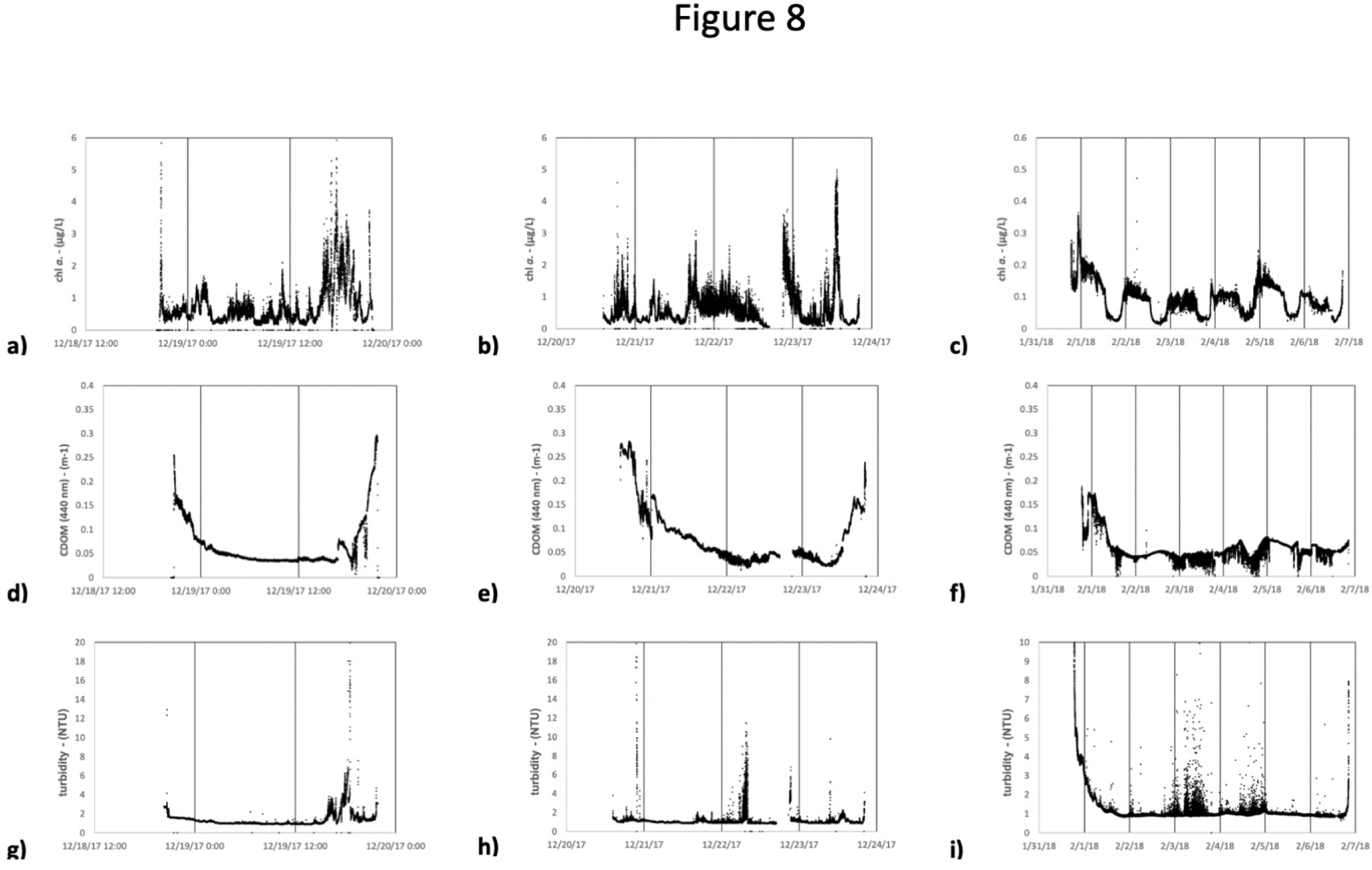
For the Dec. 18 to 19, 2017, Dec. 20 to 23, 2017, and Jan. 31 to Feb. 6, 2018 deployments, the chl. *a* time series is represented in (**A**) – (**C**) respectively, CDOM measured via fluorometric proxy is represented in (**D**) – (**F**) (explanation in text), and turbidity is represented in (**G**) – (**I**). Note, the y-axis magnitudes are different for the Jan. 31 to Feb. 6, 2018 deployment.

## 4 Discussion

### 4.1 Platform functionality

Though three deployments of increasing duration, the *Nav2* autonomous sail vehicle successfully demonstrated the potential for the platform to provide mobile, unattended monitoring of the surface coastal ocean. The *Nav2* is a unique platform in that it is small enough to be deployed in coastal and inland waters, and by functioning identically to a real sailboat it can obtain high speeds and accurately navigate and map areas of interest. The deployments demonstrated the vessel is robust enough to reliably operate and survey under non-ideal sea states with winds up to 25 knots (although we have tested the vehicle in winds > 30 knots in coastal waters of New England and Washington State). Under conditions encountered in southwest Florida with winds averaging less than 4 knots, however, the boat still managed to cover 17 to 22 NM per day, and 29 NM per day with winds averaging 8 knots (**Table 2**). These winds were not necessarily directed from behind the boat; indeed, the vessel can sail into the wind via autonomous tacking, under which significant forward progress is still made at a VMG of 10 NM per day; **Figure 4**). The vessel is capable of efficiently reaching preselected (or adjusted on the fly) waypoints (**Figures 2&3**). On the other hand, during deployment and retrieval the boat can be operated manually in sailing mode, or with a thruster (**Table 2**). The thruster is particularly useful in areas of high currents or ship traffic. Using the thruster only, the *Nav2* can be used for short missions (up to 48 hours) without the sail.

We demonstrated deployments of up to one week. The vessel was operating exceptionally at the time of retrieval and could have continued longer. Indeed, *Nav2* deployments since the time of writing this report have lasted for 15+ days. Power efficiency improvements are ongoing with multi-sensor, multi-month mission lengths feasible. Approximately 1 to 5 Watts extra power is available for sensors. The power availability and length of mission will vary with solar conditions. In solar conditions typical of Florida the panels typically provide an average of 200 W hrs day^−1^. In low light conditions typical of northern latitudes in the winter, mission planning needs to be adjusted accordingly.

Deployment or retrieval of the *Nav2* is simple but exciting and can be achieved from a boat ramp or from the beach under calm seas by a single operator. All three deployments described herein were initiated from the beach. For deployment, the operator simply walks the small hand-held trailer into the surf zone into waist-deep water until it is floating, and then slides the trailer out from underneath the vessel. The operator can leave the iOS device on shore during the actual deployment or place it into a waterproof case and hold with a lanyard. For retrieval the vehicle can be lifted back onto its wheel cart by hand in shallow water and then pulled on shore. The *Nav2* is also capable of being lifted from the water directly from a small boat. An easily overlooked aspect of using the vehicle is the attention that it garners from beachgoers. This is an opportunity for community outreach, and the southwest Florida HAB monitoring deployments were met with great inquiry and enthusiasm, eventually becoming the subject of several media features. Unfortunately, however, this curiosity also led to mission interruption on Dec. 22, 2017, when a recreational boater pulled the Nav2 from the water and proceeded towards shore until seeing the contact information and statement on the ASV. Under extremely low winds if the vessel is not obviously making forward progress it can appear “lost”. Of course, theft is always an issue, especially of a smaller 2 m length boat. Future versions of the vehicle are expected to be slightly larger to hold a larger number of sensors; this may also serve the dual purpose of being a theft deterrent. The “curiosity” effect has been better managed since these deployments by adding a large bold statement directly on the sail indicating the Nav2 is a “RESEARCH VESSEL” “TRACKED AND MONITORED AT ALL TIMES”. Boaters are increasingly aware that drones of all types on land or sea are carefully monitored. No problems with curiosity or theft have occurred since. Other ongoing improvements with the *Nav2* vehicle include an increased vehicle size to more easily accommodate a variety of sensors, the integration and testing of additional sensors, refinements to the autonomous steering algorithm to reduce oversteering and increase average speed, improved consistency of performing desirable straight data collection transects in variable currents., and improved power efficiency to provide greater power for sensors and in low light conditions.

### 4.2 Applications for Marine HAB monitoring

The utility of the platform was demonstrated for the specific application of harmful algal bloom monitoring of the Florida Red Tide species *Karenia brevis*. The recurring *K. brevis* blooms ravaging southwest Florida are challenging to monitor because blooms are most detrimental nearshore, but in many cases are transported shoreward from deeper waters ((Vargo, 2009)). While depth-resolved measurements are ideal and have been routinely obtained by glider as part of the State of Florida monitoring program, gliders have difficulty operating in waters less than 10 m deep, especially in dynamic environments. Gliders are also more expensive to operate in shallow waters (they require more frequent attention and battery and buoyancy pump servicing), prohibiting continuous operation. Finally, gliders possess a limited selection of sensors and face many sensor design constraints, currently limiting the wide-use of species-level detection techniques. Regardless, by the time *K. brevis* blooms approach the coast, they are usually at the surface of a well-mixed water column and the need for depth-resolved measurements is decreased (Robbins et al., 2006). Thus, there will for the foreseeable future be a niche that must be filled for sustained coastal surface monitoring for this species.

While the work presented herein only used a chl. *a* sensor (a common glider sensor), results serve as justification for the investment into compatible HAB species-specific sensors. The fluorescence response of organic matter has been extensively used as a proxy since terrestrially-based coastal CDOM can, for discrete regions and time intervals, display nearly linear relationships with salinity and FDOM (Coble, 1996; Del Castillo et al., 2000a). The success of the platform/sensor combination is demonstrated by the matchup to satellite observations (**Figure 6**), repeatable diel variations (**Figure 7**), obtainment of reasonable ancillary fluorometric data (**Figure 8**), and a lack of discernible bubble artifacts (**Figure 5**). Interestingly, the chl. *a* data obtained in situ was of much lower concentration than that detected by satellite. These variations can be expected based on the different nature of the measurements. The fluorometric measurements of chl. *a* can be subject to various packaging effects, especially at higher concentrations, and numerous accessory pigments can also contribute to the signal (Babin et al., 1996; Schofield et al., 2006). Satellite measurements, on the other hand, integrate over a depth interval and calculated concentrations are therefore representative of an average concentration of the surface water column. They are also more challenging in turbid and CDOM-rich optically complex waters where we conducted the deployments (Hu et al., 2005).

With a fleet of *Nav2* vehicles traveling ~20 NM per day in a repeatable triangular or “lawnmower” raster pattern, several vehicles have the potential to continuously survey a large area, regardless of water depth. The *Nav2* can also reveal more *K. brevis* surface heterogeneity than remote sensing data, and data is acquired at a much greater temporal resolution. Therefore, the *Nav2*/fluorometer package has the potential to provide satellite remote sensing ground-truthing data that can be used to improve the species-specific algorithms. An unexpected result were the repeatable diel variations observed during the longer mission (**Figure 7c**). Similar results have been observed in the same region of the surface ocean during glider missions (unpublished). While not consistent with the behavior of *K. brevis* cells, which instead exhibit positive phototaxis during the daylight hours (Schofield et al., 2006), the resident phytoplankton appear to be either altering their surface expression of chl. *a* in an excess of light or are descending to deeper waters, e.g. to obtain nutrients or alleviate photo stress (Vargo, 2009). Overall, the high sensitivity and reproducibility of the measurements highlight the functionality of the sensor for high precision measurements.

### 4.3 Other monitoring applications

While chl. *a* measurements were the primary focus of this project, the ancillary fluorometric data streams also shed light on some in water processes and allude to future applications of the *Nav2* vessel. CDOM is of interest to biogeochemists for its role in dominating ocean color, playing a critical role in photobiology, photochemistry (Helms et al., 2008), and photoproduction of CO_2_ (Clark et al., 2004), contributing to aspects of the oceanic sulfur cycle (Gali et al., 2016), and controlling the absorption of light energy and the subsequent impacts on heat flux (Hill, 2008) and other ocean-climate interactions, and in serving as a tracer of freshwater (Fichot and Benner, 2012). The fluorometrically measured CDOM exhibited intensities and spatial concentration distributions that are expected in southwest Florida (Del Castillo et al., 2000a). Earlier in the project, we did install a conductivity-temperature-depth CTD package onto the vehicle. However, conductivity measurements were unreasonable, likely due to bubble retention in the flow cell. While we still aim to resolve this issue with a different installation configuration, we can instead use CDOM as a rough proxy for salinity, with the assumption that there is a single source of freshwater input that has a high CDOM concentration (i.e. the Charlotte Harbor and the Caloosahatchee River).

The *Nav2* is inherently a meteorological sensor (e.g. for wind speed and magnitude, atmospheric temperature, and humidity). Previously, the *Nav2* has been successfully configured with fisheries sensors, including a pinger tracking hydrophone system (Sonotronics) and a cetacean and noise monitoring hydrophone (Song Meter). Trials demonstrated successful location of crab tracking pingers on the Washington coast and acoustic detection of various cetacean species. As of the time of writing, we are currently adding a Wetlabs BB3 Scatterometer and a Solinst CT logger for HABs surveys on Lake Okeechobee and the Indian River Lagoon in Florida. Addition of Oxygen/Temp Optode (Aanderaa AADI) and CT sensors as well as an ADCP (Nortec) are under consideration to provide a complete water quality monitoring suite.

## 5 Conclusions

The Navocean autonomous sail vehicle (*Nav2*) has been demonstrated to serve as a reliable mobile platform for wide-area surface coastal monitoring. To our knowledge, this is the first demonstration of a sail-driven vessel used for coastal HAB monitoring. The scientific results were shown to be reasonable and have the potential to map HAB blooms and associated environmental conditions. While the Nav2 does not capture depth variations or collect instantaneous large surface area measurements as do underwater gliders and satellites, respectively, the platform is a useful tool in the arsenal for coastal or inland monitoring. The primary benefits of using the *Nav2* vehicle are that it is fast and has reliable, autonomous navigation, has a completely renewable power source with no consumables, can function in shallow or deep water inland or offshore, and is operable by a single person. There are several additional demonstrated payload options as well as some currently in preparation. At least with the planar-style optical sensors, bubbles do not appear to contribute significant artifacts.

Harmful cyanobacterial blooms are increasing in intensity in global freshwater bodies (Paerl et al., 2018). The *Nav2* vehicle is ideal for monitoring blooms in these frequently shallow lakes, especially by limnologists who may have less training with more traditional oceanographic tools. To this end, we are readying for deployments for freshwater *Microcystis aeruginosa* HAB monitoring in Lake Okeechobee in the winter 2018-2019. Until the summer of 2018, there were few traditional monitoring efforts, and no real-time water quality monitoring sensors on Lake Okeechobee, and even now, only one stationary optical sensor is providing ground-truthing data for satellite efforts. We plan to augment this fixed location monitoring with *Nav2* surveys to both add a mobile monitoring element, but also to constrain the spatial variability of the surface optical properties in relation to remote sensing data. A second 3-channel fluorometer is currently being installed to provide phycocyanin and phycoerythrin measurements that help discriminate multiple algal species. Eventually, we envision the *Nav2* platform as an essential part of multiple monitoring programs.

## 6 Acknowledgements

We would like to thank L. Kellie Dixon, Jim Hillier, and Karl Henderson at Mote, and last but not least our former high school intern Gabriel Rey for assistance with field work.

## 8 Author Contributions

JB authored the manuscript, provided scientific oversight, and participated in field campaigns, EA is the lead designer of the autonomous vehicle hardware and software, RC developed live data visualization interface, EM coordinated field campaigns, and SD designed the sailboat and conducted deployments.

## 10 Data Availability Statement

All datasets generated for this study are included in the manuscript and the supplementary files.

## Footnotes

1 https://www.windfinder.com/windstatistics/southwest_of_tampa_bay_buoy

